# Flexible Asexuality: Naturally occurring variation in mechanisms of parthenogenesis within lineages and individuals of a facultative parthenogen, *Megacrania batesii*

**DOI:** 10.64898/2026.03.30.715418

**Authors:** Soleille M. Miller, Daniela Wilner, Jigmidmaa Boldbaatar, Lee Ann Rollins, Russell Bonduriansky

## Abstract

Parthenogenesis is relatively rare and often regarded as an evolutionary dead end. Despite this, certain parthenogenetic animal species have endured for millions of years, but it is unclear what enables the persistence of some parthenogenetic lineages. Transitions from sexual to parthenogenetic reproduction can occur through different evolutionary processes that give rise to diverse cytological reproductive mechanisms. These mechanisms are likely to influence genetic diversity, especially in the early stages after the transition to parthenogenesis and may thus affect lineage persistence. To understand such evolutionary transitions, we used experimental crosses to investigate the mechanism of parthenogenesis and the immediate genetic consequences of switching from sexual to parthenogenetic reproduction in the facultatively parthenogenetic phasmid *Megacrania batesii*. We obtained DNA sequence data from multiple lineages propagated over three generations via sex, parthenogenesis, or transitions between reproductive modes. We quantified heterozygosity and within-family genetic variation and compared the genetic patterns with predictions for known mechanisms of parthenogenesis. We found that a single generation of parthenogenesis typically resulted in (near-)complete loss of heterozygosity and an absence of within-family genetic variation, consistent with automixis with gamete duplication or terminal fusion and little/no recombination. However, we also found evidence of variation in the mechanism of parthenogenesis among lineages and even within the same individual, associated with drastic differences in the amount of heterozygosity and within-family genetic variation maintained across generations. Our findings show that considerable variation in parthenogenetic mechanisms can exist within populations and suggest that such variation could influence the persistence and evolution of parthenogenetic lineages.

## Introduction

In theory, parthenogenetic reproduction – the development of unfertilized eggs – should be favoured by selection, at least in the short term. This is because parthenogenetic females can reproduce twice as quickly as their sexual counterparts, as they do not allocate resources to the production of males, who are incapable of reproduction themselves (the two-fold cost of males; Smith 1978). Though this advantage of parthenogenesis has been demonstrated experimentally (Gibson, Delph, and Lively 2017) sexual reproduction is, paradoxically, the dominant reproductive mode in animals (Bell 1982; Otto 2009).

Nonetheless, parthenogenesis has proven a successful strategy for some lineages, helping them to survive, prosper, and even evolve for millions of years (Judson and Normark 1996). To understand why parthenogens sometimes persist but sometimes suffer extinction, a fruitful approach is to investigate how parthenogenetic reproduction affects genetic diversity and evolvability, both in the short-term and the long-term (see Bast et al. 2018; Tvedte, Logsdon, and Forbes 2019; Danchin et al. 2011). However, while a number of studies have compared ancient sexual and parthenogenetic species, few studies have investigated the immediate consequences following transitions to parthenogenetic reproduction for genetic diversity at different levels (i.e. individual, family, population).

Transitions to parthenogenesis can happen suddenly, driven by factors such as spontaneous mutations, hybridization events, or endosymbionts (reviewed in Simon et al. 2003), which can result in differing mechanisms though which parthenogens reproduce. In fact, a variety of cytological mechanisms are utilized by parthenogenetic animals, yielding different genetic and evolutionary outcomes (Suomalainen et al. 1987). Because the main advantage of sex over parthenogenesis is thought to be the greater genotypic diversity, potential for genetic purging, and evolvability enabled by sexual reproduction (Muller 1932,1964; Felsenstein 1974; Bell 1982; Agrawal 2006; Keightley and Otto 2006), the consequences of various mechanisms of parthenogenesis for genetic variation might be especially relevant to understanding the fate of newly established parthenogenetic lineages. Thus, the specific mechanism of parthenogenesis could help explain why certain parthenogenetic lineages persist in nature while others do not.

Two widespread cytological mechanisms of parthenogenesis result in unreduced chromosome sets in the offspring: apomictic (mitotic) and automictic (meiotic) parthenogenesis. In apomixis, organisms do not go through meiosis and genetic recombination, thus yielding offspring that are genetically identical to their parent (save for de novo mutations). This preserves the parental ratio of heterozygous to homozygous loci (Suomalainen et al. 1987; Pearcy et al. 2006). By contrast, in automixis, parthenogenetic eggs go through the early stages of meiosis similarly to eggs in sexual species (chromosomes pair at zygotene, crossing over occurs, followed by chromosome reduction into four azygoid nuclei), but the last stage of meiosis is altered to restore diploidy in the parthenogenetic offspring. After meiosis II, amphimixis—the fusion of the maternally produced egg cell with a paternally produced sperm during fertilization—is replaced by automixis—the fusion of two maternally produced meiotic products (i.e., the egg cell and one of the azygoid polar bodies). Diploidy in automixis can be restored through multiple mechanisms—central fusion, terminal fusion, and gamete duplication—which, in turn, result in differing genetic outcomes in the offspring (Stenberg and Saura 2009; see Fig. 2).

In central fusion, the egg cell nucleus fuses with one of its non-sister azygoid polar body nuclei following the completion of meiosis II, giving rise to the diploid embryo (see Fig. 2). In the absence of recombination between a locus and its associated centromere, the parental allelic state is maintained, including the heterozygosity at that locus. However, when recombination occurs, parental heterozygosity can be eroded, leading to offspring that are homozygous at that locus (Fig. 2b). Overall, this results in a roughly 0-33% loss of heterozygosity between the parent and the offspring; the rate at which heterozygosity erodes is dependent on the number of chiasmata formed between the locus and the centromere (Pearcy et al. 2006). This mechanism is found in many species, including the African honeybee *Apis mellifera* (Verma and Ruttner 1983), the crustacean *Artemia parthenogenetica* (Nougué et al. 2015), and the bdelloid rotifer *Adineta vaga* (Simion et al. 2020).

Automixis with terminal fusion involves the same early stages of meiosis as central fusion, but after meiosis II, the egg cell nucleus fuses with its sister azygoid polar body nucleus (see Fig. 1). Terminal fusion causes complete homozygosity in offspring except at loci that recombine with the centromere (Asher 1970). This results in a 30-100% loss of heterozygosity, depending on the amount of recombination occurring between loci and their associated centromeres (Pearcy et al. 2006). Terminal fusion appears to occur in the stick insect *Extatosoma tiaratum* (Alavi et al. 2018), and during cases of occasional parthenogenesis in some vertebrates (Lampert 2008).

**Figure 1:**
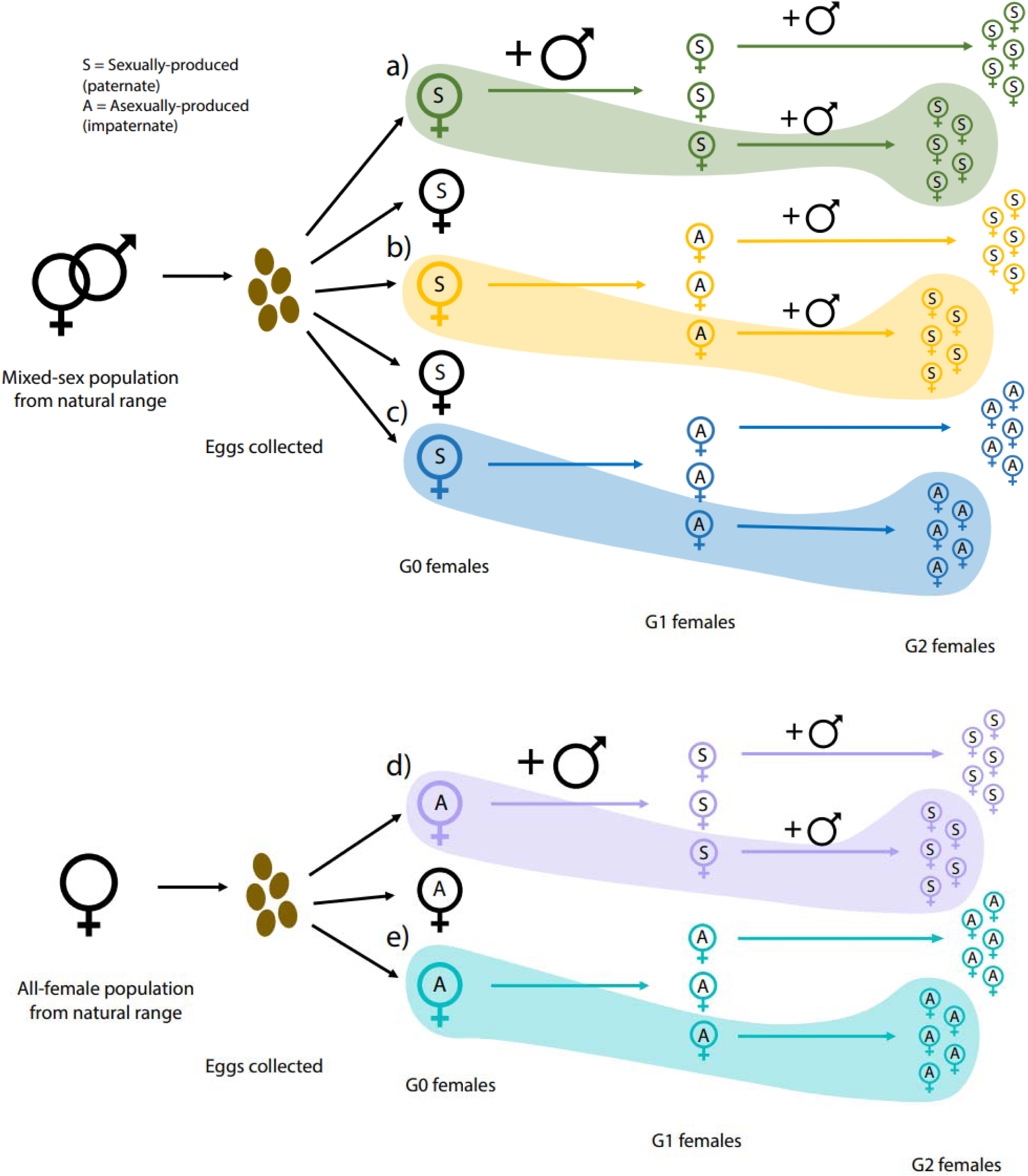
Experimental crosses and reproductive pathways. Eggs were collected from natural mixed-sex and all-female populations, reared to adulthood in the laboratory, and then allocated to one of five reproductive pathways involving sexual (“S”) and/or asexual/parthenogenetic (“A”) reproduction: SSS (a), SAS (b), SAA (c), ASS (d), and AAA (e). Colours of G0 female symbols indicate distinct matrilines (G0 females and their G1 and G2 descendants), whereas shaded areas indicate distinct lineages (G0 and G2 females that are either a mother or daughter of a unique G1 female). The “S” or “A” inside each female symbol represents whether that individual was produced sexually (paternate) or asexually/parthenogenetically (impaternate).

In gamete duplication, the egg cell nucleus duplicates itself and the two resulting nuclei then fuse together to restore diploidy (see Fig. 1). Regardless of crossing over during meiosis I, this results in complete homozygosity, except for de novo mutation. If the mother is heterozygous, gamete duplication can result in parthenogenetic offspring that are completely homozygous but genetically different from one another (i.e. clonal diversity). This is because offspring could inherit either of the homologous chromosomes (with different haplotypes) from the heterozygotic parent (Stenberg and Saura 2009). Gamete duplication happens in the termite *Cavitermes tuberosus* (Fournier 2016), the stick insect *Bacillus rossius* (Pijnacker 1969), and can be induced by the bacterial endosymbiont *Wolbachia* in *Trichogramma* wasps (Stouthamer and Kazmer 1994). Thus, the wide range of cytological mechanisms utilized by parthenogenetic organisms can lead to varied outcomes in offspring heterozygosity, which can directly affect fitness through effects on inbreeding depression (Charlesworth and Charlesworth 1987; Acevedo-Whitehouse et al. 2003; Reid et al. 2007; Chapman et al. 2009; Neaves et al. 2015), as well as mutation load and evolvability (Muller 1964; Felsenstein 1974; Lynch et al. 1995; Keightley and Otto 2006). It is therefore crucial to correctly identify the specific mechanism of parthenogenesis in use when drawing comparisons between sexual and parthenogenetic lineages.

Stick and leaf insects of the order *Phasmatodea* (phasmids) display many different mechanisms of parthenogenesis (Pijnacker 1967, 1969; Bedford 1978; Scali 2009; Schwander and Crespi 2009,2025; Alavi et al. 2018). There is also extensive chromosomal variation across the order, even including variation within species (Craddock 1972). Some phasmid species can flexibly switch between sexual and parthenogenetic reproduction—a phenomenon known as facultative parthenogenesis. In these species, females can reproduce sexually when mated and parthenogenetically when they do not mate. This makes them particularly useful model systems when comparing the consequences of sexual reproduction versus parthenogenesis: both reproductive modes can be observed within the same species, thus eliminating potential species-specific variables that might contribute to observed differences.

*Megacrania batesii* is a facultatively parthenogenetic stick insect that is being used to investigate the ecology and evolution of sexual versus parthenogenetic reproduction in both lab (Wilner et al. 2025) and natural populations (Boldbaatar et al. 2024; Miller et al. 2024; Miller 2025).The mechanism of parthenogenesis employed by *M. batesii* is unknown, but extreme homozygosity in natural parthenogenetically reproducing populations suggests that parthenogenesis involves some form of automixis (Miller et al. 2024). In the wild, *M. batesii* populations show a geographic pattern of differing reproductive modes, with some populations having an approximately equal ratio of males to females and typically exhibiting sexual reproduction (hereafter referred to as “mixed-sex” populations) but other populations having only females and exhibiting exclusively parthenogenetic reproduction (hereafter referred to as “all-female” populations). Reduced-representation whole-genome sequencing revealed that some individuals in all-female populations displayed elevated levels of heterozygosity relative to most other individuals from the same populations, despite no males having been recorded in those populations over 6 years of sampling and that the heterozygous individuals showed no evidence of genetic admixture from mixed-sex populations (Miller et al. 2024). This difference in heterozygosities could result from variation in the mechanism of parthenogenesis. Indeed, some parthenogens have been shown to display multiple mechanisms of parthenogenesis within a species (Suomalainen et al. 1987). Miller et al. (2024) also showed that there were multiple genetically distinct parthenogenetically reproducing lineages with very little within-lineage genetic diversity across *M. batesii’s* natural range. It is not known whether differences in genotype between these distinct lineages are associated with variation in the parthenogenetic mechanisms they employ.

Here, we investigated the mechanism of parthenogenesis in *M. batesii* and its consequences for genetic variation by estimating changes in heterozygosity and within-family genetic variation across generations. We first generated detailed predictions on how each known cytological mechanism would be expected to affect heterozygosity and genetic differentiation over one and two generations of parthenogenetic reproduction. We then genotyped individuals sourced from experimental lineages established from natural populations and propagated parthenogenetically, sexually, or via transitions between sexual and parthenogenetic reproduction over two generations in the laboratory. We quantified within- and between-lineage loss of heterozygosity across generations as well as genetic differentiation within families, and then we compared the observed patterns with the predictions. We first considered the mechanism observed in a large majority of lineages and then described apparent deviations from this pattern.

## Methods

### Study system and design

The Peppermint Stick Insect, *Megacrania batesii* is endemic to the wet tropics of far-north Queensland, Australia (Miller et al. 2024; Cermak and Hasenpusch 2000). Females of this species can flexibly switch between sexual and parthenogenetic reproduction (i.e. facultative parthenogenesis). When *M. batesii* females mate, some or all of their eggs can be fertilized, and fertilized eggs develop with equal probability into male or female offspring that have a father (“paternate”). Females descended from long-established all-female populations are resistant to fertilization and produce few or no paternate offspring after mating (resulting in highly female-biased offspring sex ratios, since males develop only from fertilized eggs), whereas females descended from non-resistant populations produce mostly paternate offspring and near-even sex ratios (Wilner et al. 2025). When *M. batesii* females reproduce without a male (or lay unfertilized eggs after mating), the unfertilized eggs hatch into exclusively female offspring that have no father (“impaternate”) (Wilner et al. 2025). The genotypic data used in this study were obtained from *M. batesii* samples from experimental lineages established using eggs collected from natural populations and propagated over two generations (Wilner et al. 2025). Briefly, eggs (G0) were collected from several natural mixed-sex and all-female populations within *M. batesii’s* range in far-north Queensland, Australia (see Miller et al. 2024). These eggs were reared in the lab and, when the stick insects became adults, the females were either paired with a male or housed individually to reproduce parthenogenetically. A subset of their offspring (G1) was then reared to adulthood in the lab and adult G1 females were either paired with a male or housed alone to reproduce parthenogenetically, generating G2 broods (see Fig. 1). We obtained DNA samples from G0, G1 and G2 individuals that had been frozen at −80°C (as adults or hatchling nymphs) for the analyses described below.

We sequenced DNA of females from four different types of experimental lineages (hereafter, “reproductive pathways”). To determine the rate of heterozygosity loss over one and two generations of parthenogenesis, we sampled paternate G0 females that originated (as eggs) from natural mixed-sex populations (see Miller et al. 2024) and then propagated these G0 females and their G1 daughters parthenogenetically (without males) in the laboratory to create first-generation (G1) and second-generation (G2) impaternate females (SAA reproductive pathway; see Fig. 1c). To determine how much heterozygosity is maintained after more than 3 generations of parthenogenesis, we sampled impaternate G0 females (as eggs) from natural all-female populations (see Miller et al. 2024) and propagated these G0 females and their G1 daughters parthenogenetically (AAA reproductive pathway, see Fig. 1e). To investigate the maintenance of heterozygosity through sexual reproduction, we sampled paternate G0 females (as eggs) from mixed-sex populations and propagated these G0 females and their G1 daughters sexually by pairing them with males (SSS reproductive pathway, see Fig. 1a). Lastly, to determine whether heterozygosity would be completely restored in a parthenogenetic lineage after one generation of sexual reproduction, we sampled paternate G0 females that originated (as eggs) from mixed-sex populations in the wild, propagated these G0 females parthenogenetically to produce impaternate G1 females, and then propagated those G1 females sexually (SAS reproductive pathway; see Fig. 1b).

There is high genetic differentiation between both mixed-sex and all-female populations of *M. batesii* in the wild (Miller et al. 2024), so we took eggs (G0 samples) from 4 genetically distinct mixed-sex populations and 5 genetically distinct all-female populations throughout the range to determine whether genetic populations varied in the mechanism of parthenogenesis. We also sampled multiple lineages descended from individual G0 females to investigate within-population variation. We hereafter refer to all individuals descended from an individual G0 female as a “matriline” (Fig. 1 colour) and refer to all descendants of a G0 female through one of her G1 daughters as a “lineage” (Fig. 1 shading). Our total data set consists of 35 matrilines, 59 lineages, and 555 genotyped individuals.

### Sequencing

DNA extraction and purification was conducted using the Gentra Puregene Tissue Kit (4g) from Qiagen according to the manufacturer’s protocol, “DNA Purification from Mouse Tail Tissue Using the Gentra Puregene Mouse Tail Kit,” with slight modifications to suit the study species (see Miller et al. 2024). Purified DNA was then sent to Diversity Arrays Technology Pty. Ltd. (Canberra, Australia) for targeted genotyping using the DArTag^TM^ service. DArTag^TM^ allows for the genotyping of a large number of individuals at a subset of variable loci across the genome (Guppy et al. 2018, 2020). The targeted DArTag^TM^ markers used in this study were derived from a panel of 12,977 markers across the genome created using a restriction enzyme double digest of PstI and HpaII and optimized for *M. batesii* by DArT standard pipelines (Miller et al. 2024; Kilian et al. 2012).

### Quality control filtering

Quality parameters including call rate, read depth, and polymorphic information content (PIC), was calculated using the DArTsoft analytical pipeline (Kilian et al. 2012). The custom DArTag panel initially consisted of 300 targeted single-nucleotide polymorphisms (SNPs). Before quality control filtering, the mean call rate was 0.98, and 95% of SNPs had a call rate above 96%. Quality control filtering was conducted using dartR (Gruber et al 2018) using various filtering parameters. SNP sequences matching the following parameters were removed from the dataset: locus call rate < 95% and read depths lower than 10 and greater than 175 (to remove loci in potentially duplicated regions; see Fig. S1). Additionally, we removed individual samples that were missing data for more than 50% of markers. This quality control filtering removed 40 SNPs (13%) and 6 individual samples (1%). After quality control, 260 high quality SNP markers were retained for further analysis in 555 sequenced *M. batesii* females.

### Mechanism Predictions

We identified specific predictions following one and two generations of parthenogenesis under 6 different cytological diploidy-restoration mechanisms: gamete duplication with/without recombination, terminal fusion with/without recombination, and central fusion with/without recombination. These predictions were based on single-generation genetic predictions in Suomalainen et al (1987) and extrapolated to a second generation of parthenogenesis. All-female *M. batesii* populations in the wild had much lower heterozygosity than did mixed-sex populations (Miller et al. 2024), so we can reject a priori most mechanisms that result in genetic outcomes similar to apomixis (i.e., complete maintenance of heterozygosity), like gonoid thelytoky or premeiotic doubling and subsequent reduction. However, we derived predictions for automixis via central fusion with no recombination (which has the same genetic outcomes as apomixis) to compare with the other mechanisms.

We first outline the predictions that best differentiate the various mechanisms and represent these predictions visually in figures 2 and 3. Under gamete duplication, we would expect to see a complete loss of heterozygosity in one generation of parthenogenesis regardless of recombination (Fig. 2). If G1 mothers had differing genotypes (assuming no recombination), they would be highly differentiated (See Fig. 3a). Moreover, the G0 paternate mothers would be genetically different from their G1 impaternate offspring, but G1 impaternate mothers would be genetically identical to their G2 daughters, and G2 daughters within the same lineage would be genetically identical to each other (Fig. 3a). The only prediction that changes if one recombination event is assumed to occur is that there would be higher potential for genotype variation among G1 offspring of the same matriline (Fig. 3b). This would result in the largest potential genetic differentiation across G1 individuals within matrilines of the three mechanisms. In other words, this mechanism is expected to generate the lowest correlation between genotype matrices of G1 siblings within the same matriline when compared to the other mechanisms.

**Figure 2:**
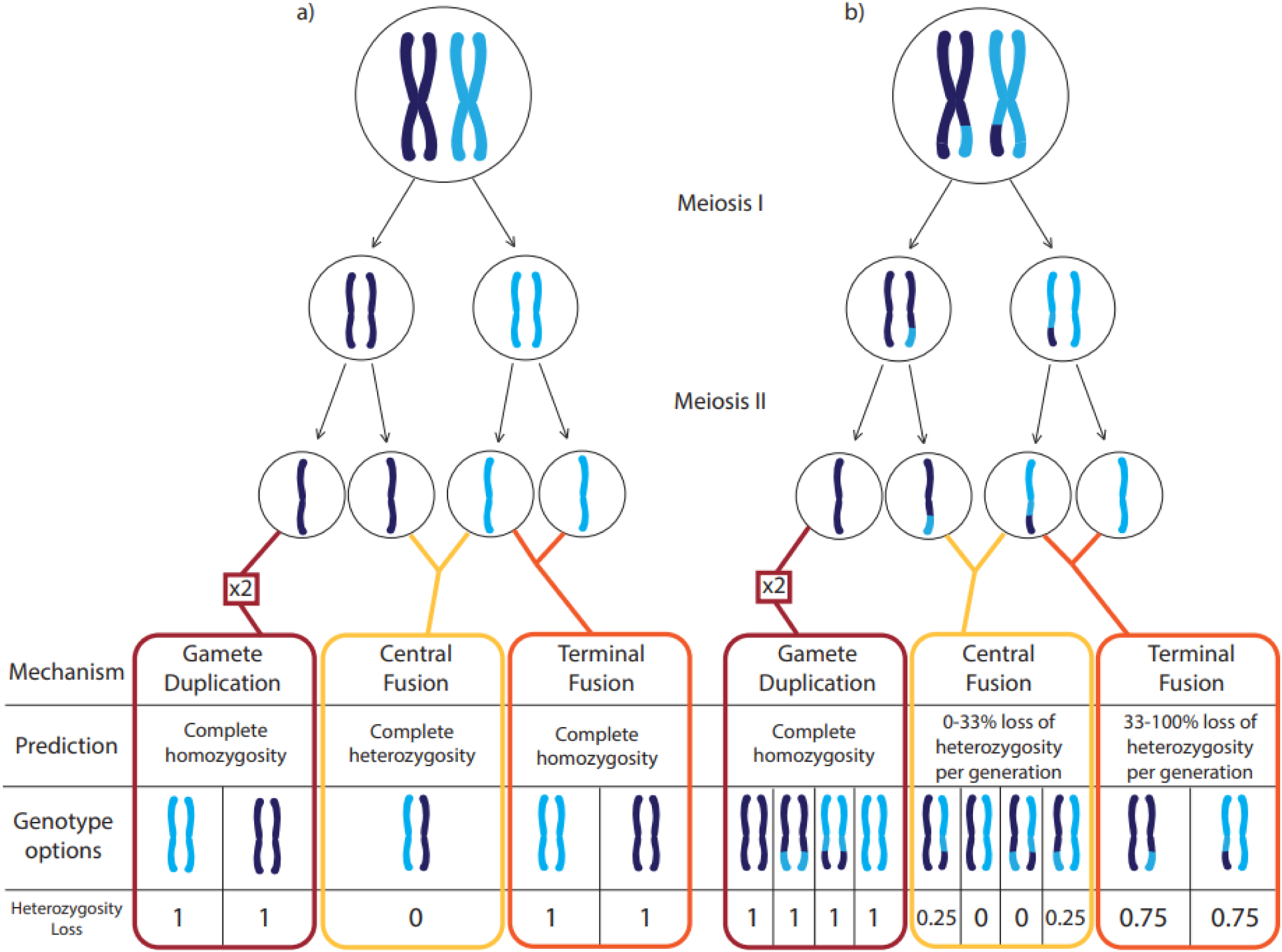
The three main diploidy-restoration mechanisms and their heterozygosity outcomes (as proportion heterozygosity loss) in automictic parthenogenesis over a single generation: Each mechanism allows for recombination, and we therefore represent the genetic outcomes after one generation of parthenogenesis if these mechanisms occur a) without recombination or b) with one recombination event. The box showing “x2” represents the gamete duplicating itself and fusing back together to restore diploidy (gamete duplication).

**Figure 3:**
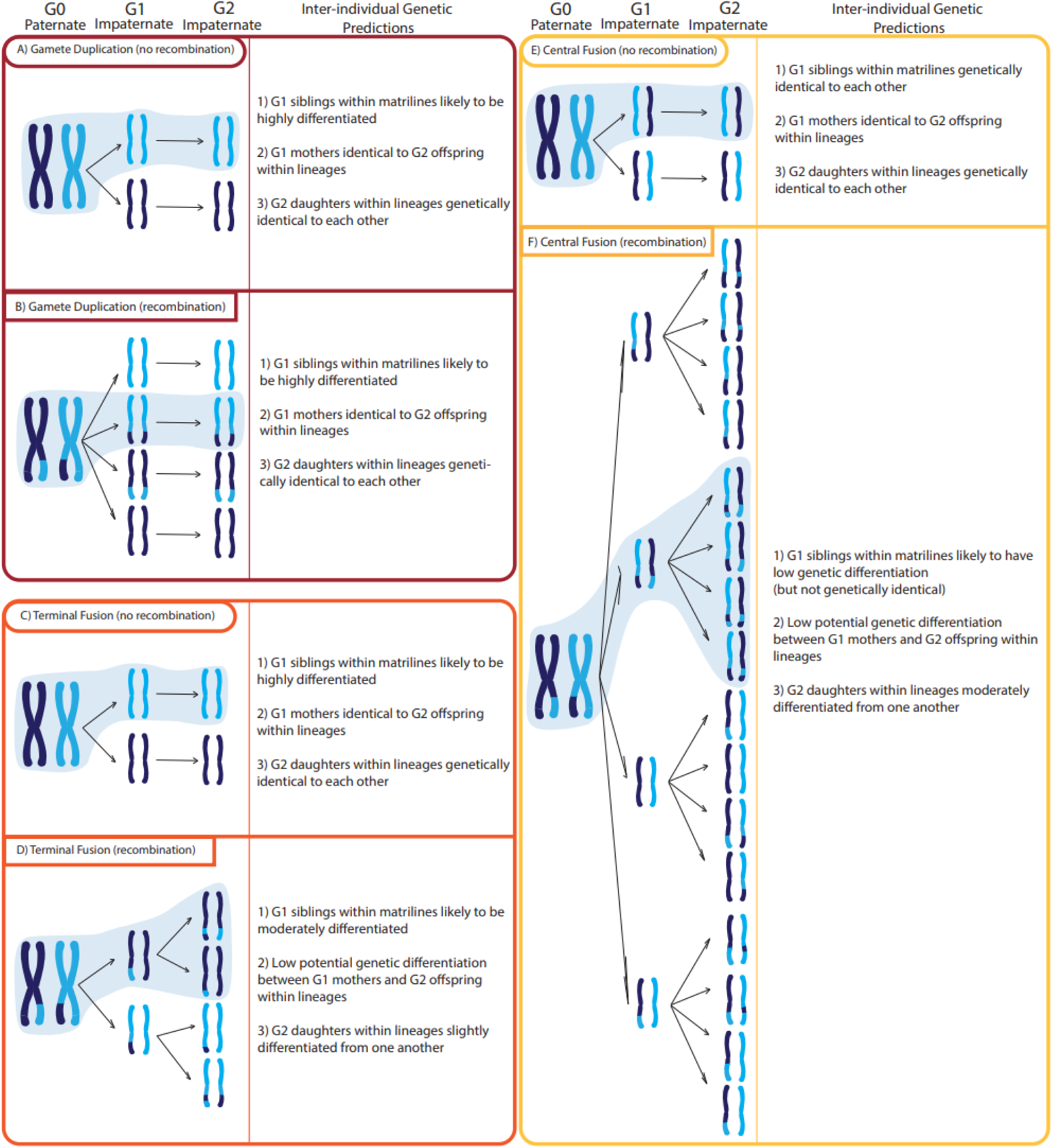
Predicted within-family genetic variation after one and two generations of automictic parthenogenesis via gamete duplication a) without recombination and b) with one recombination event, terminal fusion c) without recombination and d) with one recombination event, and central fusion e) without recombination and f) with one recombination event. This assumes that the same cytological mechanism is happening in both the first and second generation of parthenogenesis. G0 paternate represents the G0 individuals sourced from wild mixed-sex populations, G1 impaternate represents the first generation of parthenogenesis, and G2 impaternate represents the second generation of parthenogenesis. Inter-individual genetic predictions are the predictions based on the possible genotypes for each mechanism

Under terminal fusion with no recombination, we would expect to see heterozygosity and genetic differentiation patterns that are the same as gamete duplication with no recombination (see Fig. 2 and Fig. 3c). Similarly to gamete duplication, we would expect that G0 mothers would be genetically different from their G1 offspring, G1 mothers would be genetically identical to their G2 daughters, and G2 daughters within the same lineage would be genetically identical to each other (Fig. 3c). However, assuming one recombination event during automixis with terminal fusion, we would expect to see a 33-100% loss of heterozygosity each generation, with loci only maintaining heterozygosity when they have recombined (Fig. 2). There would still be low potential G1 genotype variation and high G1 genetic differentiation within matrilines, but G1 mothers would be slightly genetically differentiated from their G2 offspring. Additionally, recombination would increase genotypic variation within and genetic differentiation between G2 offspring within a lineage (see Fig. 3d).

Central fusion with no recombination results in predictions similar to apomixis: all individuals in the matriline would be genetically identical clones of each other (excluding de novo mutations) and there would be complete maintenance of heterozygosity across all generations (see Fig. 2 and Fig. 3e). Lastly, in central fusion with recombination, we would expect to see a 0-33% loss of heterozygosity with each generation of parthenogenesis, with recombining loci becoming homozygous (Fig. 2). This mechanism has the potential to produce a greater number of different G1 genotypes per mother, but those genotypes would be less genetically differentiated than under terminal fusion or gamete duplication (see Fig. 3f). There would be low potential genetic differentiation between G1 mothers and G2 offspring, with a small chance that G0, G1, and G2 individuals of the same lineage are genetically identical. Central fusion with recombination would also result in lineages with the most potential G2 genotype variation and genetic differentiation between G2 siblings when compared to the other mechanisms (Fig. 3f).

### Analysis

We first sought to identify the predominant mechanism of parthenogenesis in *M. batesii*. We then investigated the individuals that showed deviations from this mechanism, hereafter referred to as variants.

To determine the predominant mechanism, we first excluded the lineages that had apparent variants (see below). Then, we calculated the percentage of heterozygosity lost in one generation of parthenogenesis using the remaining 83 paternate mother-impaternate daughter pairs (pooling G0-G1 and G1-G2 pairs). To do this, we identified the loci that were heterozygous in paternate mothers for each mother-daughter pair (both G0 and G1 paternate mothers) and calculated the loss of heterozygosity at those loci in their impaternate (parthenogenetic) offspring (both G1 and G2) using a custom R script (supplementary material “analysis.qmd”). For this analysis, we only included loci that were heterozygous in the mothers to see how heterozygosity was lost for each locus, instead of comparing observed heterozygosity rates. We chose to do this as it is a more informative and direct way to measure the loss of heterozygosity (Jaron et al. 2022).

To see how heterozygosity changed after two generations of parthenogenesis, we took lineages that had reproduced parthenogenetically twice (SAA reproductive pathway) and calculated the loss of heterozygosity between the first and second generation of parthenogenesis, only using loci that were also heterozygous in the G0 female of that matriline. This analysis was also done using the same custom R script (supplementary material “analysis.qmd”) using *tidyverse* (Wickham et al. 2019) and *dartR* (Gruber et al. 2018). To see if there was a loss of heterozygosity between generation G0 and G1 and generation G1 and G2, we ran respective pairwise Wilcoxon signed-rank tests using the *wilcox.test()* function in base R.

To compare the level of heterozygosity between recently parthenogenetic lineages and lineages that have been reproducing parthenogenetically for many generations, and determine if a single generation of parthenogenesis is enough to reduce heterozygosity values to the same level, we also calculated heterozygosity rates of AAA lineages (i.e., lineages that descended from a parthenogenetic G0 ancestor sourced from a natural all-female population and then reproduced parthenogenetically twice in the laboratory). We then compared G1 individuals from the SAA reproductive pathway to G1 individuals from the AAA reproductive pathway using an independent-samples Wilcoxon signed-rank test. There was little variation in the number of heterozygous loci across parthenogenetic generations (See Fig. 4c; sd=1.94), so we used the average number of heterozygous loci across the three generations as the “parthenogenetic baseline”. Likewise, we estimated changes in heterozygosity across generations in lineages that descended from a paternate G0 ancestor sourced from a natural mixed-sex population and then reproduced sexually twice in the laboratory (SSS reproductive pathway, sexual baseline; Fig. 4c).

**Figure 4:**
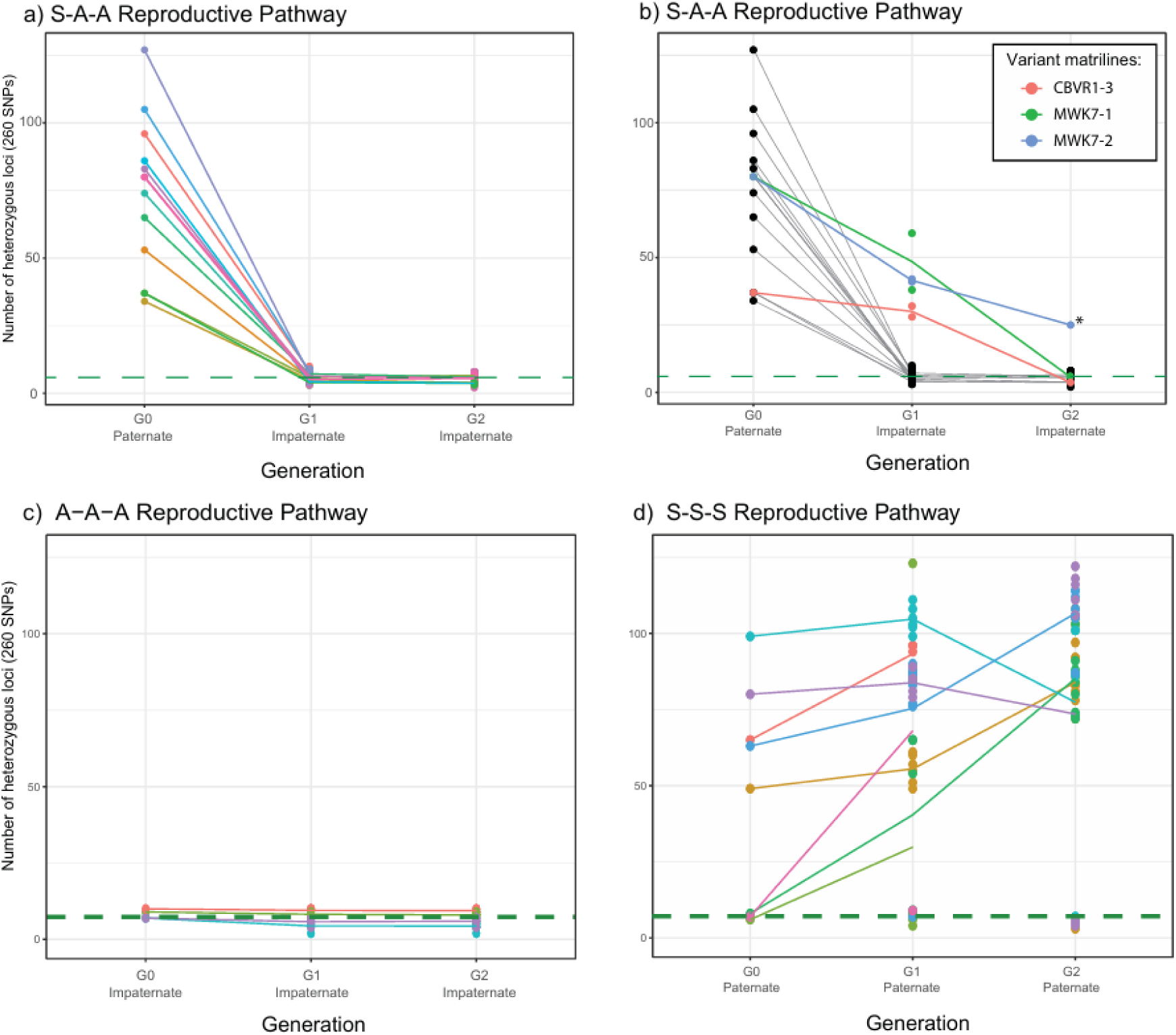
Reproductive pathways and resulting heterozygosity. (A&B) The number of heterozygous loci (out of 260) after paternate G0 females reproduced parthenogenetically and their impaternate daughters also reproduced parthenogenetically (SAA pathway). Variant lineages were excluded from (a) but included and highlighted in (b). Loci that were already homozygous in G0 females were excluded from the count for G1 and G2. (C and D) (C) individuals taken from all-female populations in the wild and all of their impaternate descendants (AAA pathway) and (D) individuals taken from mixed-sex populations and all of their paternate descendants (SSS pathway). Colour represents matriline identity and points represent the average number of heterozygous loci in each generation for each matriline. The parthenogenetic baseline as identified in (C) is included here as a dashed green line.

A small number of loci were identified as heterozygous in the AAA reproductive pathway, and this number showed little change across generations (see Results). To determine whether these loci are a random subset of all loci (as would be expected if they were due to sequencing error), or whether specific loci maintain heterozygosity within matrilines and thus represent genuine inherited genetic diversity, we tested whether the heterozygous loci within matrilines were more correlated to their matriline than to a randomly assigned matriline using a randomization test. In this test, we first recoded each locus in every individual as either heterozygous or homozygous. Then we randomly constructed simulated matrilines from among the sampled individuals, keeping the sample size of each constructed matriline the same as that of the genuine matriline. We then created correlation matrices for each constructed matriline and calculated the mean correlation among individuals in that matrix. We ran this over 5,000 iterations and plotted the distribution of mean correlations for all simulated matrilines (Fig. S2). Lastly, we calculated the 95% confidence interval of this distribution and compared it with the mean observed correlation of individuals to their actual matrilines.

A few females obtained as eggs from mixed-sex populations, or produced by male-paired females in the lab, were found to be impaternate (see Results), and these individuals and their descendants were excluded from further analyses involving the SSS and SAA reproductive pathways. To see if one generation of sexual reproduction restores heterozygosity in parthenogenetic lineages to the sexual baseline (SSS), we calculated heterozygosity rates of lineages that originated from a mixed-sex population, reproduced parthenogenetically, and then were mated and reproduced sexually (SAS reproductive pathway).

After estimating the degree of loss of heterozygosity each generation, we used *pvclust()* (Suzuki and Shimodaira 2006) to visualize the genetic relationships between individuals in each matriline (including all individuals showing the predominant mechanism and the mechanism variants) to compare with predictions. For this cluster analysis, we included all 260 high quality loci. We calculated a dissimilarity matrix of the samples using the *bitwise.dist()* function in the R package “poppr” (Kamvar et al. 2014) based on Hamming distances (see Wang et al. 2015). This matrix was then used with *pvclust()* to create a dendrogram based on correlation distance between individuals, and 10,000 bootstrap iterations were conducted to determine the level of support for each node (Suzuki and Shimodaira 2006).

For direct comparisons of genetic differentiation between G1 individuals within matrilines (for inter-individual genetic prediction #1; see Fig. 3), we first calculated the correlation matrix of genotypes of G1 individuals within matrilines using the cor() function. We then omitted any loci that were missing a genotype call in any of the samples within that matriline. We then found the mean and standard deviation of the genetic correlations between individuals within each matriline using the *mean()* function from base R and the *std()* function from the package *pracma* (Borchers 2025). We did the same analysis to quantify genetic differentiation between G2 individuals within the same lineage. This was done for lineages showing the predominant mechanism as well as in lineages that were apparent variations to the predominant mechanism (2 daughters from matriline MWK7-1, 2 daughters from MWK7-2, and 2 daughters from CBVR1-3) and again with G2 individuals from lineages descended from these G1 variants.

## Results

### Heterozygosity in Parthenogenetic vs. Sexual Matrilines

In lineages that were sourced from natural all-female populations and underwent a further 3 generations of parthenogenesis in the laboratory (AAA reproductive pathway), low levels of heterozygosity were observed throughout all generations. The average proportion of heterozygous loci in these lineages was 6.8 (sd = 2.1) out of 260 (hereafter referred to as the “parthenogenetic baseline”; see Fig. 4c). By contrast, in lineages sourced from mixed-sex populations that underwent 3 generations of sexual reproduction in the laboratory (SSS reproductive pathway), the average number of heterozygous loci was 88.9 (sd = 18.4).

When following lineages through an SAS pathway (see Fig. 1), we found that heterozygosity could be restored in one generation of sex to similar levels of heterozygosity (Fig. S3).

When analysing the SSS reproductive pathway, we found that 11 of 89 sequenced individuals (G0=2, G1=4, G2=5, derived from 6 different matrilines) had very low heterozygosity values indicative of parthenogenetic reproduction. This shows that *M. batesii* females produce some parthenogenetic offspring despite mating, and that females in mixed-sex populations in the wild occasionally reproduce parthenogenetically despite the availability of males. These impaternate individuals and their descendants were excluded from any analyses involving the SSS, SAA, and SAS reproductive pathways.

### Predominant Mechanism

In most cases, parthenogenetic descendants of paternate *M. batesii* females exhibited a very large (mean = 90.2%, se = 0.43) decrease in observed heterozygosity levels over one generation (G0-G1; Wilcoxon paired-samples signed-rank test: p<0.001) and 91.3% (se = 0.55) decrease over two generations (G0-G2; n=77 and n=54 respectively; see Fig. 4a) relative to the paternate G0 ancestor. The value of the initial loss of heterozygosity is consistent with predictions for automixis with terminal fusion and small amounts of recombination (Fig. 3 & Table 1). However, the average decrease in heterozygosity between the first and second generation of parthenogenesis was only 5.6% (se=0.63) or 0.33 loci (G1-G2; n=54), with 74% of second-generation parthenogens showing no decrease in heterozygosity relative to their impaternate mothers. This pattern is not consistent with predictions for any form of automixis, as all mechanisms are expected to result in a continuous decrease in heterozygosity with each generation until complete homozygosity is reached. Instead, we found that most impaternate G1 individuals were at the parthenogenetic baseline level of heterozygosity after only one generation of parthenogenesis. Indeed, when we compare G1 individuals deriving from paternate mothers (through the SAA reproductive pathway; mean number of heterozygous loci=6.8, sd=0.17) to G1 individuals deriving from impaternate mothers (through the AAA reproductive pathway; mean number of heterozygous loci=7.7, se=0.26), we find no difference in mean heterozygosity (Wilcoxon independent-samples signed-rank test: p=0.938). This suggests that heterozygosity reaches a stable minimum level after a single generation of parthenogenesis by a paternate female (Fig. 4a). Interestingly, the loci that appear to retain heterozygosity following parthenogenetic reproduction are matriline-specific and therefore do not appear to be random genotyping errors (Fig. S4; see Discussion).

**Table 1:**
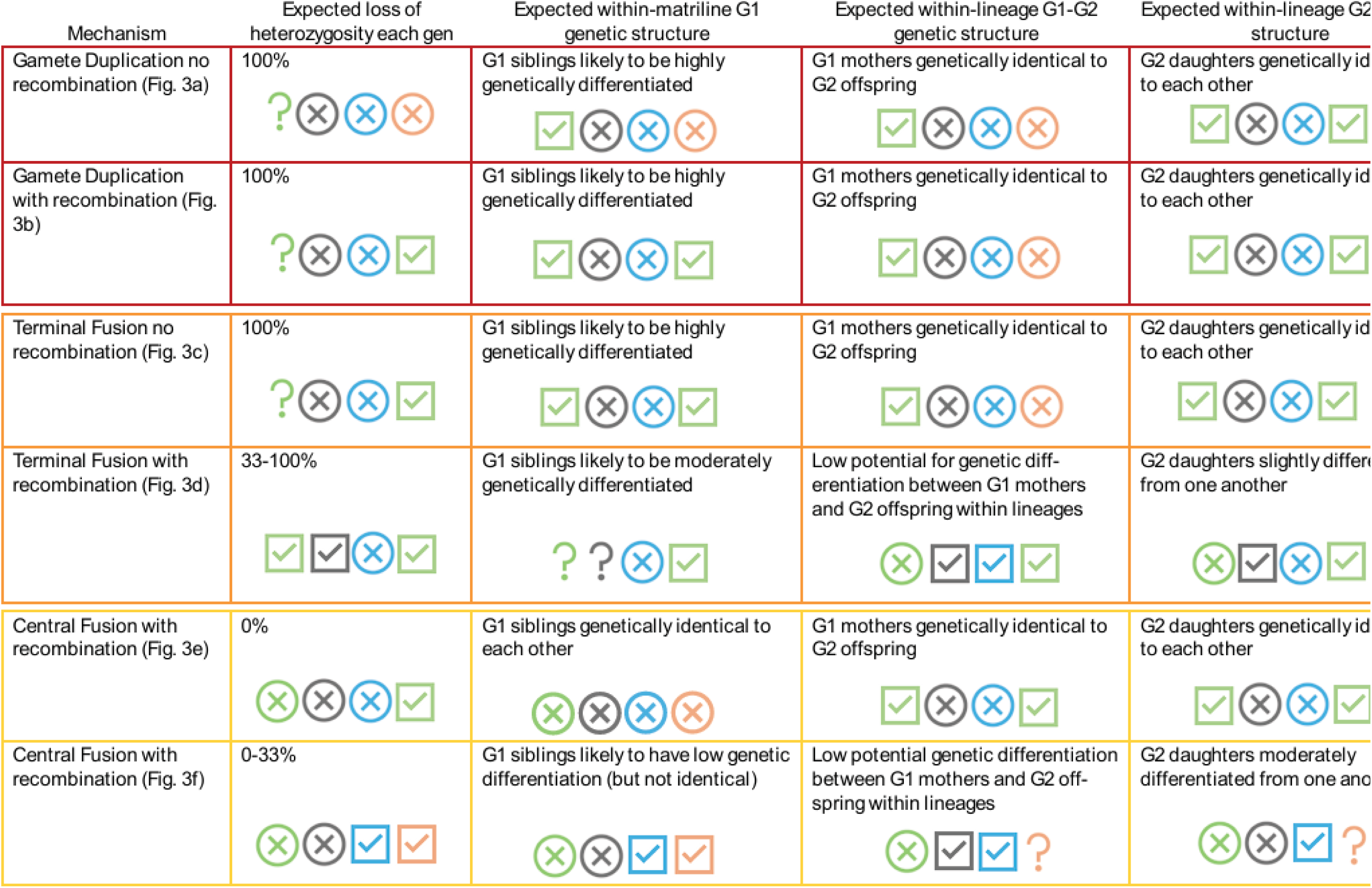
Predictions and results for heterozygosity loss and genotype diversity after one and two generations of parthenogenesis (starting from a paternate female ancestor) under 3 different cytological mechanisms of parthenogenesis with and without recombination. Results are shown as coloured icons, with the predominant mechanism shown in green, and the variant matrilines that show apparent deviations from this mechanism shown separately: MWK7-1 (grey), MWK7-2 (blue), and CBVR (orange). Checkmark icons represent results that are consistent with a prediction, question mark icons represent results that are inconclusive (i.e. not clearly consistent with a prediction; see Discussion), and X icons represent results that are inconsistent with a prediction. The border colours of the table correspond to the colours assigned to each cytological mechanism in figures 2 and 3: Gamete duplication (dark red), Terminal fusion (bright orange), and Central fusion (gold).

| Mechanism | Expected loss of heterozygosity each gen | Expected within-matriline G1 genetic structure | Expected within-lineage G1-G2 genetic structure | Expected within-lineage G2 structure |
| --- | --- | --- | --- | --- |
| Gamete Duplication no recombination (Fig. 3a) | 100% | G1 siblings likely to be highly genetically differentiated | G1 mothers genetically identical to G2 offspring | G2 daughters genetically identical to each other |
| Gamete Duplication with recombination (Fig. 3b) | 100% | G1 siblings likely to be highly genetically differentiated | G1 mothers genetically identical to G2 offspring | G2 daughters genetically identical to each other |
| Terminal Fusion no recombination (Fig. 3c) | 100% | G1 siblings likely to be highly genetically differentiated | G1 mothers genetically identical to G2 offspring | G2 daughters genetically identical to each other |
| Terminal Fusion with recombination (Fig. 3d) | 33-100% | G1 siblings likely to be moderately genetically differentiated | Low potential for genetic differentiation between G1 mothers and G2 offspring within lineages | G2 daughters slightly different from one another |
| Central Fusion with recombination (Fig. 3e) | 0% | G1 siblings genetically identical to each other | G1 mothers genetically identical to G2 offspring | G2 daughters genetically identical to each other |
| Central Fusion with recombination (Fig. 3f) | 0-33% | G1 siblings likely to have low genetic differentiation (but not identical) | Low potential genetic differentiation between G1 mothers and G2 offspring within lineages | G2 daughters moderately differentiated from one another |

Taken together, these findings suggest that the predominant mechanism is either gamete duplication or terminal fusion with no recombination. Both of these mechanisms are thought to result in complete loss of heterozygosity after a single generation of parthenogenesis. The low levels of apparent heterozygosity that we found remaining after a single generation of parthenogenetic reproduction are therefore not expected under these mechanisms. However, this apparent heterozygosity may not represent genuine heterozygous loci (see Discussion).

Genetic cluster results show that G1 females of the same matriline (i.e., the same G0 mother) have moderate genetic differentiation from one another but appear to be genetically identical to their G2 offspring, and that all G2 siblings appear to be genetically identical to each other (see Fig. 5). We also found that G1 siblings produced with the predominant mechanism within the same matriline were, on average, more genetically differentiated from each other when compared to the G1 siblings that were considered variants to the predominant mechanism (See Table 2). These genetic differentiation patterns are consistent with either gamete duplication or terminal fusion with no recombination (Fig. 3 & Table 1).

**Figure 5:**
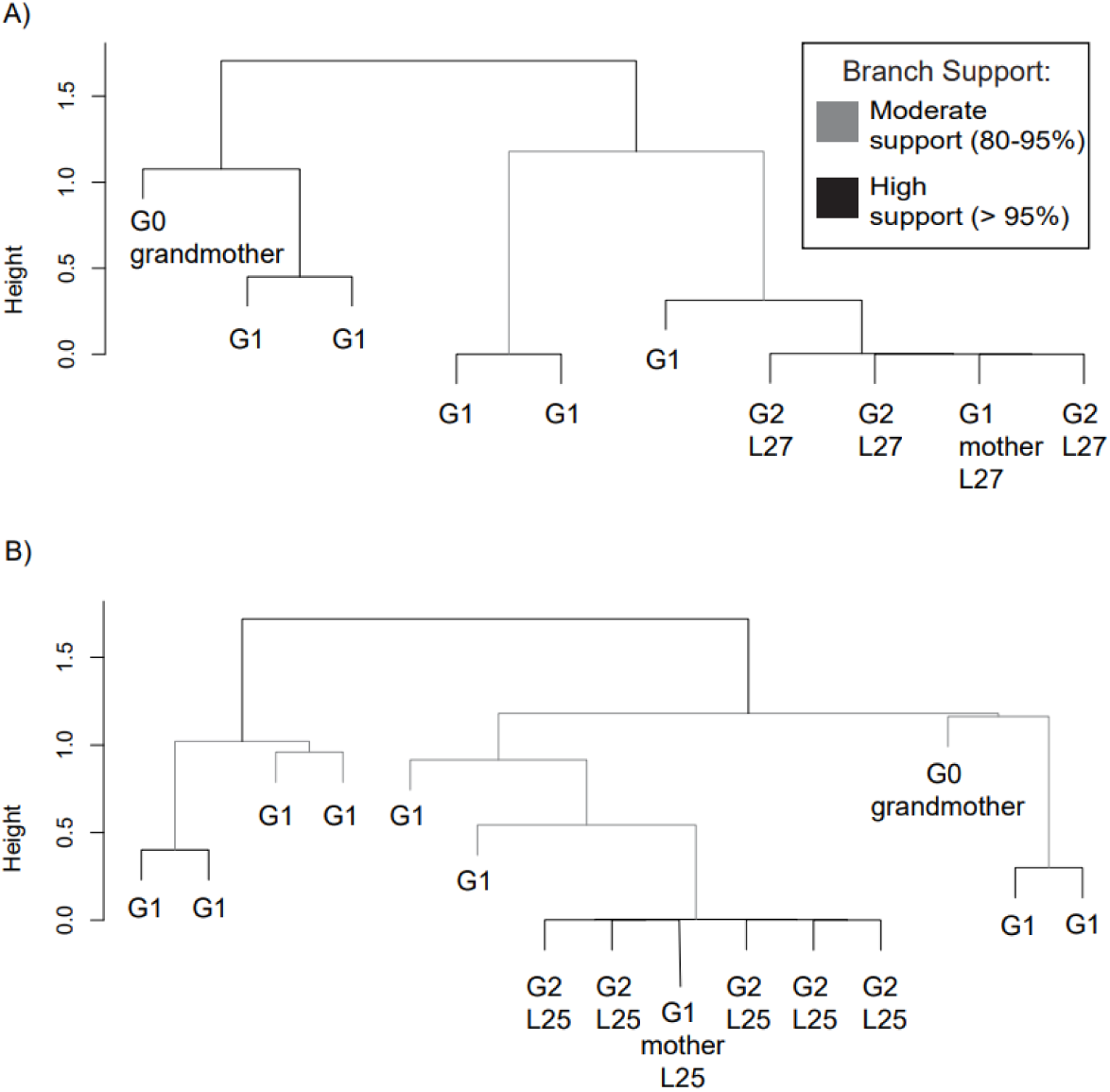
Cluster dendrograms showing genetic relationships between females within representative matrilines and lineages that exhibit the predominant mechanism of parthenogenesis: a) CBVR1-2 and b) CO2-2. These matrilines are from geographically and genetically distant populations, yet show similar patterns of within-matriline genetic structure. Branches are shaded according to the level of bootstrapped support (based on 10,000 bootstrap iterations). G0 paternate females (labelled G0 grandmother) were sourced from natural mixed-sex populations, and G1, and G2 represent descendants produced parthenogenetically in the laboratory. Unnamed G1 individuals are daughters of the G0 grandmother but did not contribute sequenced offspring to the analysis.

**Table 2:** Genetic correlations among G1 individuals within matrilines as determined by genotype calls from DArTseq data. Matrilines with * denote variant individuals that appeared to reproduce via terminal fusion with recombination and matrilines with ** indicate variant individuals that appeared to reproduce via central fusion with recombination. Some individuals had missing genotype calls; these loci were removed from this analysis. Therefore, the number of loci refers to the total number of loci analysed in that matriline out of the 260 loci that passed quality filtering. Cells with NA can be found in matrilines consisting of only two G1 individuals, as there is no standard deviation when calculating the mean correlation between just two individuals.

| Matriline | Mean correlation | Standard deviation | Number of individuals sequenced | Number of loci |
| --- | --- | --- | --- | --- |
| 1 | 0.6105 | 0.0818 | 3 | 260 |
| 4 | 0.7567 | 0.0873 | 3 | 260 |
| 16 | 0.8838 | 0.0351 | 9 | 260 |
| 17 | 0.8566 | 0.0287 | 9 | 259 |
| 18 | 0.8794 | 0.0213 | 4 | 260 |
| 19 | 0.7761 | 0.0810 | 6 | 257 |
| 22 | 0.7106 | 0.0448 | 5 | 258 |
| 24 | 0.7773 | 0.0454 | 3 | 260 |
| 25 | 0.6088 | 0.0784 | 6 | 260 |
| 28 | 0.3177 | 0.2594 | 4 | 259 |
| 29 | 0.7279 | NA | 2 | 256 |
| 31 | 0.7010 | 0.0596 | 18 | 258 |
| 32 | 0.6902 | 0.0683 | 5 | 260 |
| 18* | 0.9729 | NA | 2 | 259 |
| 31* | 0.8833 | NA | 2 | 260 |
| 32** | 0.8059 | NA | 2 | 260 |

### Mechanism Variants

While 92.8% of the mother-offspring pairs in the first generation of the SAA reproductive pathway showed similar, dramatic loss of heterozygosity over a single generation of parthenogenetic reproduction and were thus considered to be employing the same mechanism of parthenogenesis, several individuals deviated strikingly from this pattern. The heterozygosity losses in these individuals (“variants”) were 18-38 standard deviations lower than in typical (i.e., predominant mechanism) samples in the G1 generation, and 16 standard deviations lower than in typical samples in the G2 generation. Thus, the variants appear to deviate from the predominant mechanism. The variant subset included six impaternate G1 individuals and one impaternate G2 individual and showed a heterozygosity loss of 13% to 52% over one generation of parthenogenetic reproduction (see Fig. 4b & Table 1).

The seven mechanism variants came from 3 matrilines, originating from female MWK7-1 (Matriline 31), female MWK7-2 (Matriline 32), and female CBVR1-3 (Matriline 18). MWK7-1 and MWK7-2 both descended from a single female from mixed-sex population “MK” (part of the Northern genotype cluster) and CBVR1-3 descended from mixed-sex population “VR” (part of the Southern genotype cluster) (Miller et al. 2024). Three G1 variants went on to reproduce parthenogenetically again, and their second-generation impaternate offspring showed patterns consistent with the main mechanism, with one exception (denoted with an asterisk in Fig. 4b and Fig. 6a).

**Figure 6:**
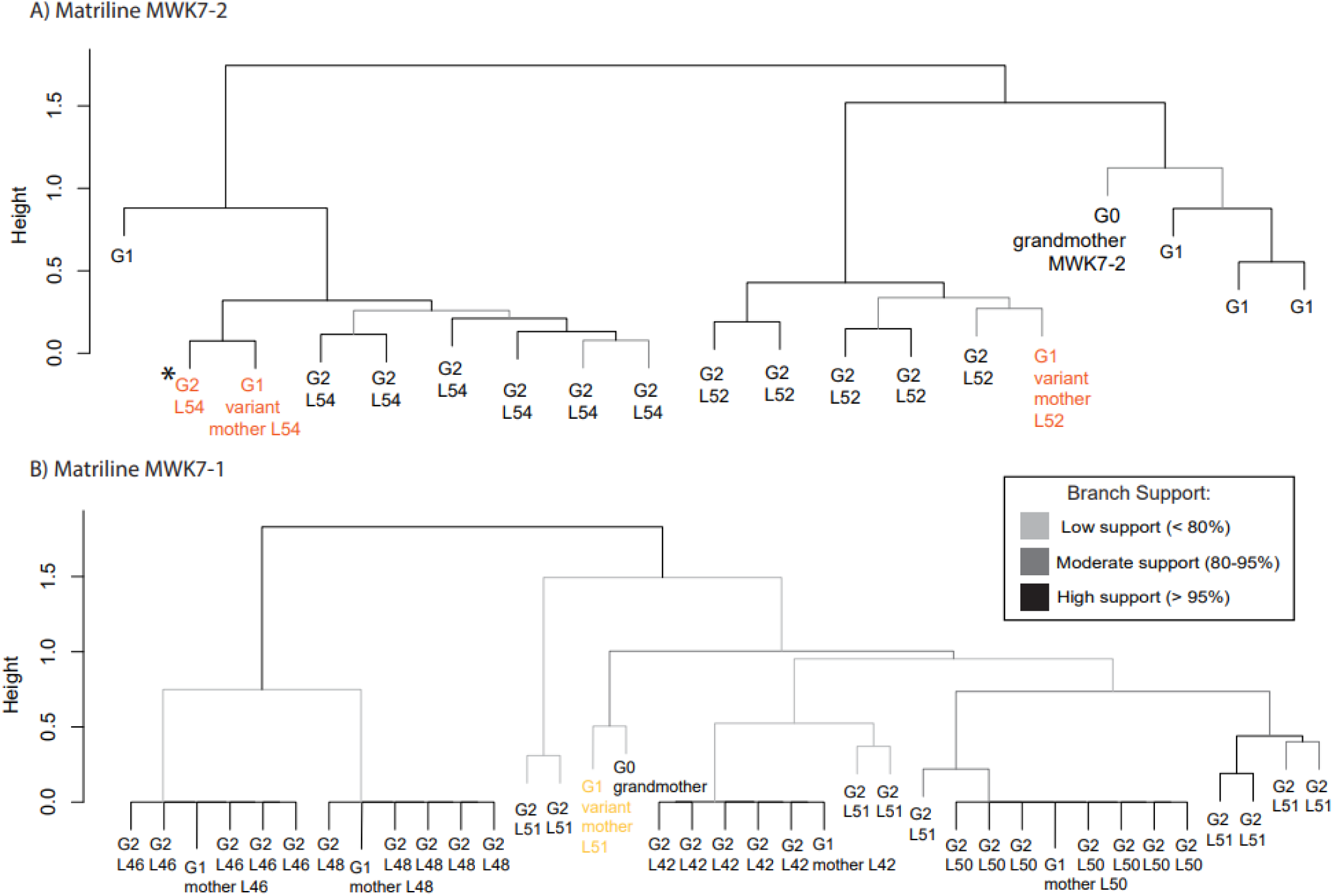
Dendrogram showing genetic relationships between individual females and lineages from the variant matrilines. A): In matriline MWK7-2, both G1 variant lineages L54 and L52 show genetic clustering patterns consistent with predictions for automixis with terminal fusion and recombination. B): In matriline MWK7-1, G1 variant mother L51 and offspring show genetic clustering patterns consistent with predictions for automixis with central fusion and recombination, but all other lineages show patterns consistent with gamete duplication or terminal fusion with little to no recombination. Branches are shaded according to the level of bootstrapped support (based on 10,000 bootstrap iterations). G0 paternate females (labelled G0 grandmother) were sourced as eggs from natural mixed-sex populations, and G1, and G2 represent the generation of parthenogenesis in the laboratory. Females that show heterozygosity losses consistent with terminal fusion and central fusion with recombination (i.e., variant mechanisms) are highlighted in orange and yellow respectively. The asterisk denotes the individual that showed relatively high heterozygosity after two generations of parthenogenesis. Unnamed G1 individuals are daughters of the G0 grandmother but did not contribute sequenced offspring to the analysis.

Two G1 variants were daughters of female CBVR1-3 (Matriline 18). They had absolute heterozygosity losses consistent with the predictions of automixis with central fusion (14% and 23% per generation) but were frozen as hatchlings and did not go on to reproduce so we do not have G2 genetic structure data to compare with our predictions. However, these variant G1 individuals had an average genetic correlation that was higher than the average for matrilines showing the predominant mechanism (Table 2). This G1 genetic structure is also consistent with predictions for central fusion, which is expected to result in low genetic differentiation of G1 siblings relative to the other three mechanisms (see Table 1).

Two other G1 variants were daughters of MWK7-2 (Matriline 32). They had a loss of heterozygosity of 48-49% and cluster analysis showed that both lineages had patterns consistent with terminal fusion with recombination (Fig. 6a): there was low (but non-zero) genetic differentiation between G2 individuals within lineages and relatively high genetic differentiation between G1 offspring within the matriline (See tables 1, 2, and 3). Additionally, one of those G1 impaternate females (G1 L54; See Fig. 6) reproduced parthenogenetically again and one of her daughters displayed a heterozygosity loss also consistent with terminal fusion with recombination (G2 L46; highlighted orange in Fig. 6).

**Table 3:** Genetic correlation among G2 individuals within lineages as determined by genotype calls from DArTseq data. Matrilines with * denote variant individuals that appeared to reproduce via terminal fusion with recombination, and matrilines with ** indicate variant individuals that appeared to reproduce via central fusion with recombination. Some individuals had missing genotype calls; these loci were removed from this analysis. Therefore, the number of loci refers to the total number of loci analysed in that matriline out of the 260 loci that passed quality filtering. Cells with NA can be found in matrilines consisting of only two individuals, as there is no standard deviation when calculating the mean correlation between just two individuals.

| Lineage | Mean correlation | Standard deviation | Number of individuals sequenced | No. loci |
| --- | --- | --- | --- | --- |
| 21 | 0.9965 | 0.0025 | 5 | 260 |
| 23 | 0.9975 | 0.0021 | 5 | 259 |
| 25 | 0.9971 | 0.0020 | 5 | 256 |
| 26 | 0.9973 | 0.0025 | 3 | 260 |
| 27 | 0.9954 | 0.0038 | 3 | 258 |
| 32 | 0.9993 | 0.0010 | 4 | 259 |
| 42 | 0.9965 | 0.0025 | 5 | 258 |
| 44 | 0.9990 | 0.0012 | 10 | 256 |
| 46 | 0.9994 | 0.0009 | 5 | 259 |
| 48 | 0.9994 | 0.0009 | 5 | 259 |
| 50 | 0.9985 | 0.0013 | 7 | 260 |
| 52* | 0.8670 | 0.0721 | 5 | 258 |
| 54* | 0.8896 | 0.0519 | 7 | 260 |
| 51** | 0.8113 | 0.0821 | 9 | 258 |

Finally, two G1 parthenogenetic variants were daughters of MWK7-1 (Matriline 31). One of these daughters went on to reproduce parthenogenetically again (Lineage 51), producing offspring that showed heterozygosity loss consistent with central fusion with recombination. This lineage also showed genetic structuring consistent with central fusion with recombination: moderately differentiated G2 daughters within the lineage, and low genetic differentiation between G1 individuals within a matriline (Fig. 6b; G2 Lineage 51; Table 1, 2, and 3). However, female MWK7-1 also had daughters that were not variants – i.e., daughters and G2 descendants with heterozygosity and genetic structure patterns consistent with the main mechanism (Fig. 6b). This single individual therefore appeared to reproduce through multiple mechanisms of parthenogenesis, resulting in varying genetic diversity and structure outcomes in descendants.

## Discussion

The cytological mechanisms underlying parthenogenesis have the potential to influence the immediate consequences following a transition from sexual to parthenogenetic reproduction. This, in turn, could shape the evolutionary trajectories of parthenogenetic lineages, potentially influencing which of these lineages persist and adapt and which go extinct. In most individuals sampled from wild sexually reproducing populations of *Megacrania batesii*, we found that parthenogenetic reproduction causes immediate declines in heterozygosity. Indeed, a single generation of parthenogenetic reproduction resulted in a level of heterozygosity in offspring that is equivalent to the heterozygosity of lineages that have been reproducing parthenogenetically for many generations. Parthenogenesis in *M. batesii* typically also results in dramatic losses of within-lineage genetic differentiation in just one generation. Together, our findings suggest that the predominant cytological mechanism of parthenogenesis in *M. batesii* across their range is either automixis with gamete duplication or terminal fusion with no recombination. However, our analysis also revealed naturally occurring variants within populations that exhibited alternative cytological mechanisms of parthenogenesis. In some variant lineages, we found genetic patterns more consistent with terminal or central fusion. We even observed evidence of multiple mechanisms of parthenogenesis employed across multiple reproductive bouts by a single individual. After two generations of parthenogenesis, nearly all individuals exhibited very low heterozygosity, but one individual still exhibited strongly elevated heterozygosity. Matrilines that showed atypical mechanisms of parthenogenesis also had increased genetic differentiation within and between lineages, which persisted after two generations. By influencing genotypic diversity, this apparent variation in the cytological mechanisms of parthenogenesis could affect lineage evolution and persistence.

Intra-individual genetic diversity metrics like heterozygosity could have important implications for the persistence of parthenogenetic lineages in nature. Greater genetic diversity can provide a wider range of traits for natural selection to act upon and enable a more flexible and rapid response to new selective regimes, facilitating adaptation to changing conditions. Heterozygosity can also be important because it shields recessive alleles from being expressed, some of which can be deleterious and lead to premature mortality and reduced reproductive success (i.e., inbreeding depression; Charlesworth and Charlesworth 1987; Stebbins and Wright 1977; Lande 1994; Lynch, Conery, and Burger 1995; Vrijenhoek 1994). Indeed, changes in heterozygosity can be directly related to changes in fitness (Chapman et al. 2009), and *M. batesii* individuals that reproduce sexually tend to have higher offspring viability, though the effect was inconsistent (Wilner et al. 2024). Although impaternate individuals (both sourced from natural populations and lab-reared) showed extremely low levels of heterozygosity, some all-female populations in the wild seem to be performing well, with high fecundity and consistently large population sizes across several years of surveys (Miller et al. 2024). The success of these all-female populations might be due to high heterozygosity in these populations when they first transitioned to parthenogenesis. High initial heterozygosity could have resulted in high genotype diversity across individuals, resulting in multiple genotypically distinct lineages (i.e., multiple lineages homozygous for different alleles). This could have enabled selection to favour the most fit genotypes and ultimately resulted in the spread of the one or few high-fitness genotypes that are currently observed in those wild all-female populations.

Regarding genetic structure of *M. batesii* parthenogenetic lineages, our study revealed differentiation between lineages, but little to no differentiation among G2 siblings within the same lineage. These findings align with our predictions for automixis involving gamete duplication or terminal fusion with minimal or no recombination. Overall, we found profound genetic differences between sexual and parthenogenetic lineages of *M. batesii* in this study. These patterns align with those observed in wild populations (Miller et al. 2024). However, the present study shows that this striking difference in genetic diversity usually emerges within a single generation of parthenogenesis.

While parthenogenesis in *M. batesii* typically resulted in nearly complete homozygosity, impaternate individuals typically retained ∼2.3% heterozygous loci regardless of the number of generations of parthenogenesis that the lineage had undergone. We observed minimal differences in heterozygosity between the first and second generation of parthenogenesis, with heterozygosity remaining unchanged in most lineages (see Fig. 4a). Not a single individual was found to be completely homozygous across the 260-locus panel. This pattern is not consistent with any known mechanism of automixis since, starting from a heterozygous background, all mechanisms are predicted to either reduce heterozygosity to zero in one generation or progressively reduce heterozygosity each generation until reaching complete homozygosity (with the exception of central fusion with no recombination), the speed of this reduction being dependent on recombination rates (Suomalainen et al. 1987; De Meeûs et al. 2007).

There are a few potential explanations for why we see these low levels of heterozygosity in the parthenogenetic *M. batesii* lineages. First, this could be due to erroneous genotype calls. However, we believe this to be unlikely as there were clear patterns in which loci maintained heterozygosity across impaternate individuals (see supp Fig. S4). These loci were heterozygous in 60 - 87% of impaternate individuals, but none were heterozygous in all individuals. Alternatively, these patterns could result from duplicated regions in the *M. batesii* genome instead of genuine heterozygous loci. The few phasmid genomes that have been sequenced and assembled show genomes that are large (∼3.5 Gb) and repeat-rich (Wu et al. 2017; Stuart et al. 2023). Although *M. batesii* lacks an assembled genome sequence, it is possible that its genome shows similar patterns. Single base-pair differences between duplicated regions would be very difficult to distinguish from heterozygosity using unmapped reads. Indeed, apparent low levels of heterozygosity have previously been observed in other stick insects as well as *Arabidopsis* plants, and in *Arabidopsis* this pattern was found to result from SNPs mapping to unrecognized duplicated regions in the genome creating spurious variant calls (Jaron et al. 2022; Jaegle et al. 2023; Larose et al. 2023). If this is the case in *M. batesii*, the predominant cytological mechanism employed by *M. batesii* is likely to be gamete duplication or terminal fusion with no recombination. However, our comparison of coverage levels between loci that maintained heterozygosity and those that lost it did not reveal any consistent differences, suggesting that another mechanism may be at play and indicating a need for further research.

It is also possible that *M. batesii* lineages retain some genuine heterozygosity across many generations of parthenogenetic reproduction. If the main mechanism utilized by *M. batesii* is terminal fusion, heterozygosity could be preserved indefinitely at loci that are closely linked to the centromere (Suomalainen et al. 1987). Therefore, the observed maintenance of heterozygosity at these loci could be attributed to selection favouring recombination at specific locations in the genome during parthenogenesis (Lichten and Goldman 1995). The African cape honeybee *A. mellifera* reproduces parthenogenetically through automixis with central fusion (Verma and Ruttner 1983). However, many *A. mellifera* lineages maintain high levels of heterozygosity over many generations (Oldroyd et al. 2011) and there is evidence to suggest that natural selection is actively maintaining heterozygosity in these lineages (Goudie et al. 2012; Goudie, Allsopp, and Oldroyd 2014). It is possible that selection favours preservation of heterozygosity at certain loci in *M. batesii*, creating the patterns we found in this study. Whether this could be due to selective processes like heterozygote advantage is yet to be determined (Fisher 1930; Fisher 1990; Crow 1999). Ultimately, whether it be through duplication events or genuine heterozygosity, these patterns still contribute to the overall genetic diversity of individuals and lineages.

As well as loci that appear to maintain heterozygosity through generations of parthenogenesis, we found that 41/83 impaternate females had at least 1 locus that “became” heterozygous (i.e. their mother was homozygous at that locus). The average number of loci that were heterozygous in impaternate offspring that were not heterozygous in their mothers was 0.855 with a standard deviation of 1.14. The only way heterozygosity could be restored at a locus under any ploidy-restoration mechanism of automixis would be via spontaneous mutations (Suomalainen et al. 1987). However, it is highly unlikely that most/all offspring would gain a mutation at the same locus. This pattern instead could be explained by variant calling errors due to restriction site polymorphisms causing allelic dropout. This sort of bias is common among targeted sequencing approaches (Luca et al. 2011; Arnold et al. 2013; Davey et al. 2013; Gautier et al. 2013). Thus, the targeted sequencing approach used for this study might be underestimating the number of true heterozygous loci in individuals.

Most of the lineages in this study dropped to the parthenogenetic baseline of heterozygosity after just one generation of parthenogenesis, but we identified six “variant” lineages from three matrilines that showed heterozygosity loss patterns more consistent with terminal fusion (33-100% loss) or central fusion (0-33% loss) with recombination. In most cases, these non-typical mechanisms only appeared to persist for one generation: by the second generation of parthenogenesis, heterozygosity in these lineages also dropped to the parthenogenetic baseline. This result suggests that there is standing genetic variation in the cytological mechanisms of parthenogenesis within natural populations of *M. batesii*. Such parthenogenetic lineages that could maintain elevated heterozygosity over two or more generations of parthenogenesis might have an evolvability advantage over lineages that lose all heterozygosity over a single parthenogenetic generation, because the variants would be able to generate more genotypic diversity for selection to act on.

Two individuals belonging to the matriline “CBVR1-3” displayed intergenerational heterozygosity-loss patterns and G1 genetic structure consistent with central fusion with recombination (i.e., a heterozygosity loss of between 0-33%). Unfortunately, these G1 individuals were euthanized as hatchlings, so we lack genetic data for G2 individuals of those lineages. The two other variant matrilines (“MWK7-1” and “MWK7-2”) were sourced from a different mixed-sex population. While they are represented as two separate matrilines in this study, the G0 founders were full siblings (hatched from eggs collected from the same wild female) and thus belonged to the same “grand matriline”. Lineages derived from matriline “MWK7-2” showed patterns indicative of terminal fusion with recombination, resulting in a 48% and 49% heterozygosity loss within a single generation, G1 siblings that were relatively highly differentiated from one another, and G2 offspring that were genetically similar but not identical to their siblings. One of the G2 females from this matriline also retained relatively high heterozygosity levels even after two generations of parthenogenesis (see Fig. 4b and 7; denoted with asterisk), showing how differences in cytological mechanisms of parthenogenesis can influence lineage genetic diversity. One lineage derived from the matriline “MWK7-1” showed patterns consistent with central fusion with recombination, resulting in an average 26% heterozygosity loss in one generation and highly differentiated G2 offspring. This mother gave rise to 6 lineages that we studied, but only one lineage seemed to reproduce via central fusion (see Fig. 6). This suggests some instability in the meiotic mechanism of parthenogenesis in “MWK7-1” that could generate a range of cytological mechanisms between successive parthenogenetic eggs of a single individual. To our knowledge, this is the first time that such within-individual variation in mechanisms of parthenogenesis has been described in nature, though suggestive patterns have been noted in *Timema* (Larose et al. 2023). Descendants of this “grand matriline” also diverged from typical *M. batesii* phenotypes in body colour, and lineages descended from “MWK7-2” had highly variable hatchling viability. A more detailed description of this matriline will be provided in a forthcoming paper.

Overall, this study shows that there is standing genetic variation in the mechanism of parthenogenesis in natural populations of *M. batesii,* which may explain the elevated heterozygosity observed in certain all-female populations despite generations of parthenogenetic reproduction, and perhaps contribute to the success of some all-female populations (Miller et al. 2024). The results suggest that some naturally occurring mechanism variants can maintain elevated heterozygosity over many generations of parthenogenesis. Within-species variation in the cytological mechanism of parthenogenesis has been reported previously in other animals, including brine shrimp (Barigozzi 1974), nematodes (Lahl et al. 2006), coccids (Nur 1971), wasps (Rössler and Debach 1973; Stille 1985), *Drosophila* (Carson et al. 1969; Markow 2013) and *Bacillus* stick insects (Marescalchi and Scali 2003; Lavanchy et al. 2024). Additionally, we found that females deriving from mixed-sex populations occasionally reproduced parthenogenetically despite having been mated. Approximately 12% of females in the SSS reproductive pathway had heterozygosity levels consistent with the parthenogenetic baseline, despite having been sourced from a mixed-sex natural population or produced by a confirmed mating in the lab. This suggests that natural populations of *M. batesii* also exhibit variation in resistance to fertilization, consistent with patterns reported by Wilner et al. (2024).

The genetic differentiation found between lineages displaying mechanism variants could have important implications for the ecology and evolution of *M. batesii*. This variation observed in parthenogenetic mechanisms led to higher levels of genetic diversity within individuals and greater differentiation between individuals than expected under the predominant mechanism. This additional genotypic variability, even if it only lasts for one or a few generations, could provide recently established parthenogenetic populations with additional variation on which selection can act. After a few generations, that variation could be whittled away thorough selection, leaving only one or a few successful genotypes in the population. This is consistent with the pattern we see in natural populations of *M. batesii*, where the all-female populations usually consist of one or two genotypes, with little (if any) intra-population diversity (Miller et al. 2024). For *M. batesii*, this suggests that wild parthenogenetic individuals may currently possess genotypes well-suited to their stable tropical habitat, selection having removed the unfit genotypes from the populations.

However, due to the decline in both within- and between-individual genetic diversity after multiple generations of parthenogenesis, such populations may prove to be less resistant to future environmental change. Our findings thus illustrate how understanding the early stages of transitions to parthenogenesis could help to explain the success of some parthenogenetic lineages over others.

### Summary

This study investigates how parthenogenesis—the development of offspring from unfertilized eggs—affects genetic diversity in the stick insect *Megacrania batesii*. The authors used experimental crosses and DNA sequencing to compare offspring produced by sexual and parthenogenetic reproduction. They found that parthenogenetic reproduction often caused a near-complete loss of genetic variation, but that the extent of this loss differed among lineages and even within individuals. These results show that mechanisms of parthenogenesis can vary within populations and suggest that such variation may help explain why some asexual lineages persist over evolutionary timescales despite reduced genetic diversity.

## Data availability Statement

All data and code for this manuscript will be uploaded to a public repository during the peer review process.

## Funding

This work was supported by the Australian Research Council Discovery Grant DP200101971 to R.B.

## Conflict of interest

The authors declare no conflict of interest.

## Acknowledgements

We thank Shay Hirani, Natalie Haryanto, Richard Benedict Jurenang, and Patrick Liao for their help with insect rearing and egg collection. We also thank Dr. Katarina Stuart for her assistance with data analysis and interpretation. Thank you to DArTseq Technologies for conducting the DArT tag sequencing and for their support in generating genetic markers.

## Author contributions

Conceptualization: S.M.M.; funding acquisition: R.B.; data collection: R.B., D.W., J.B., S.M.M.; data analysis and visualization: S.M.M., L.A.R.; writing – original draft: S.M.M.; writing – review and editing: R.B., L.A.R.

## Supplementary Materials

**Fig S1:**
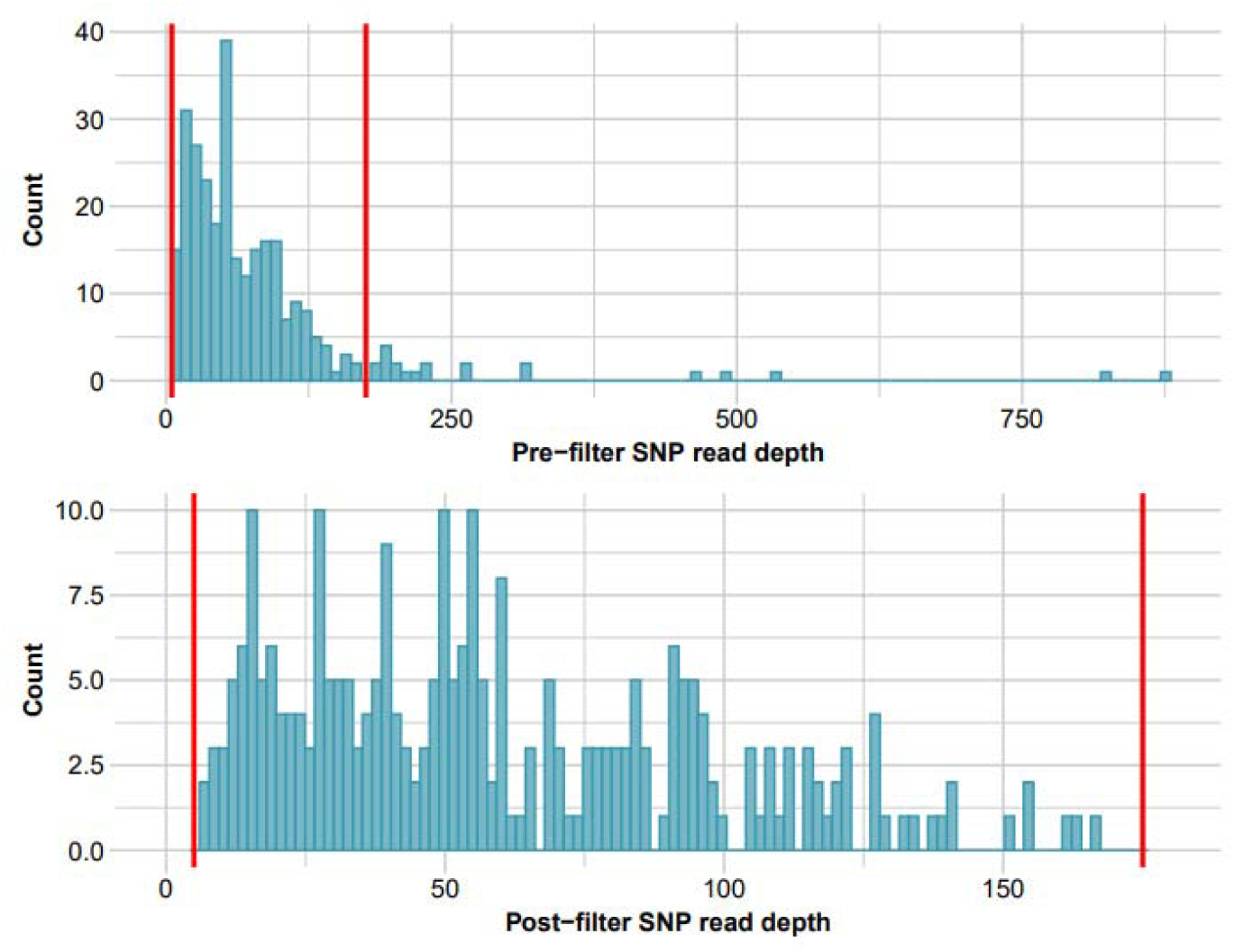
Histogram of read depths of SNPs a) before and b) after filtering for read depth greater than 3 and lower than 175.

**Figure S2:**
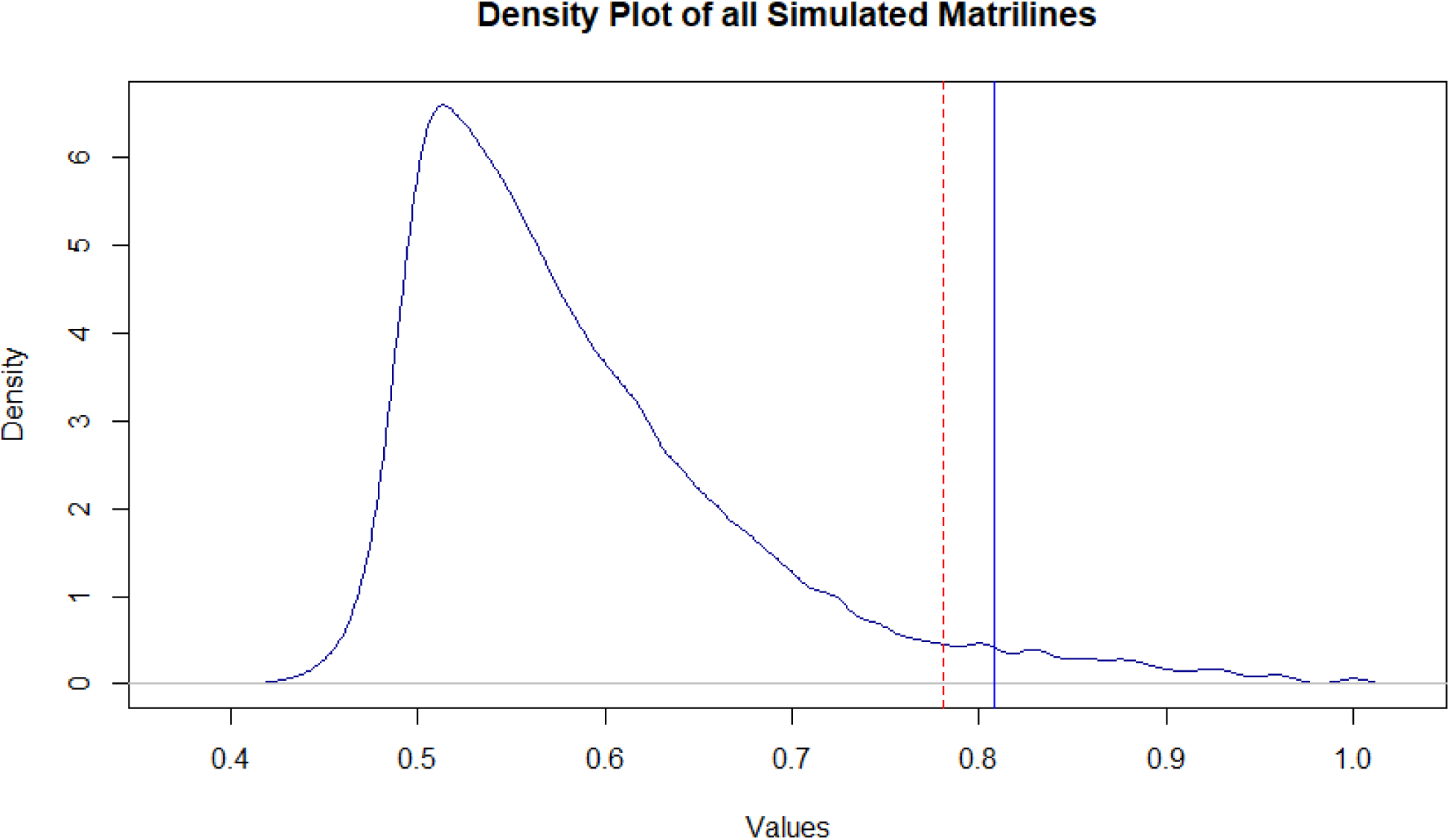
Distribution of mean genetic correlations within simulated randomized matrilines. These correlations were based off genotype calls and run for 5000 iterations of a randomization test. The red dotted line indicates the upper 95^th^ percentile in this simulated distribution and the vertical blue solid line indicates the average correlation of genuine *Megacrania batesii* matrilines.

**Figure S3:**
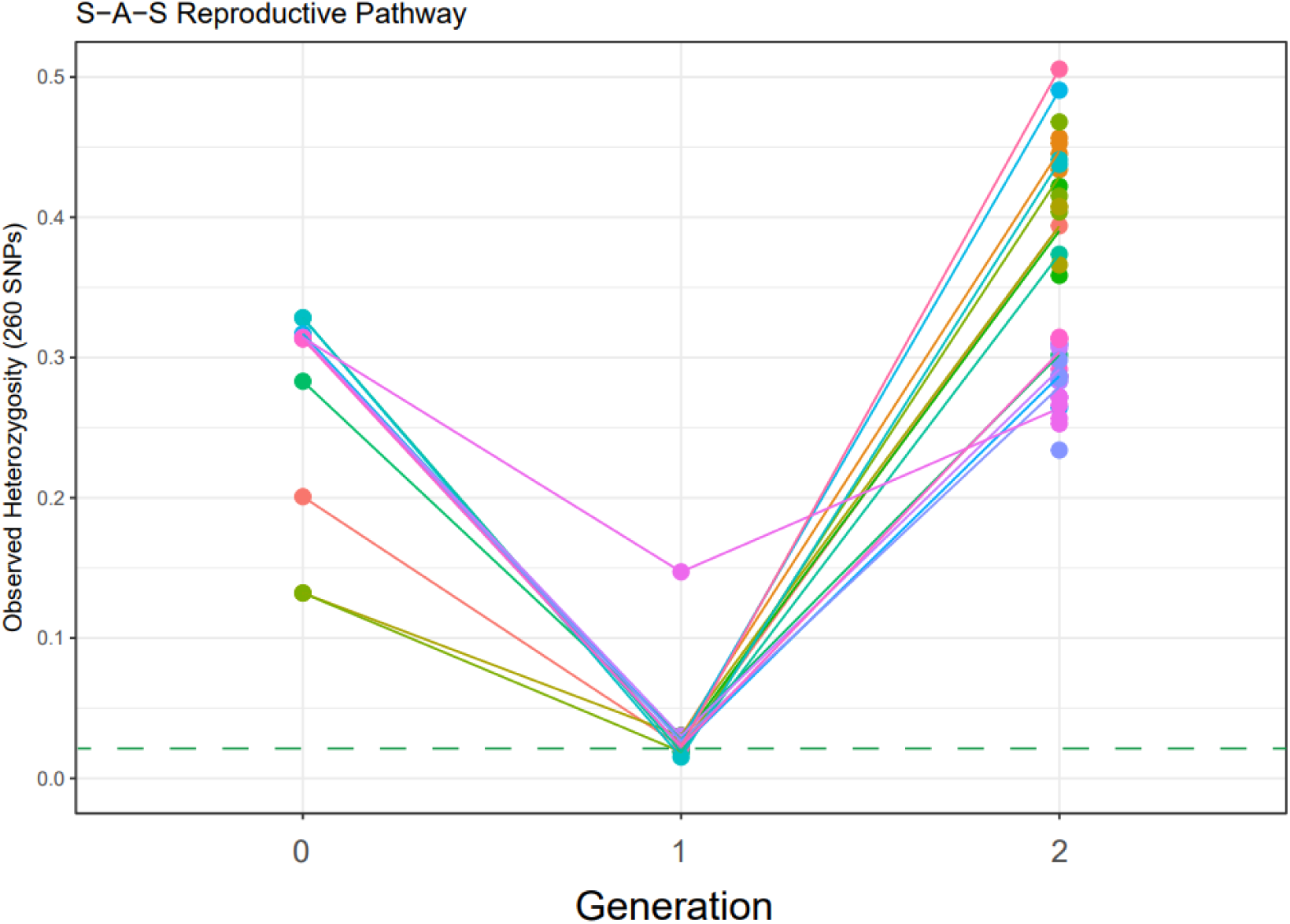
The number of heterozygous loci (out of 260) after 0 (i.e. paternate) and 1 generation of parthenogenesis, then after 1 generation (G2) of sexual reproduction (S-A-S reproductive pathway). Colour represents matriline identity and lines represent the average number of heterozygous loci in each generation for each matriline. The parthenogenetic baseline is included here as a dashed green line.

**Figure S4:**
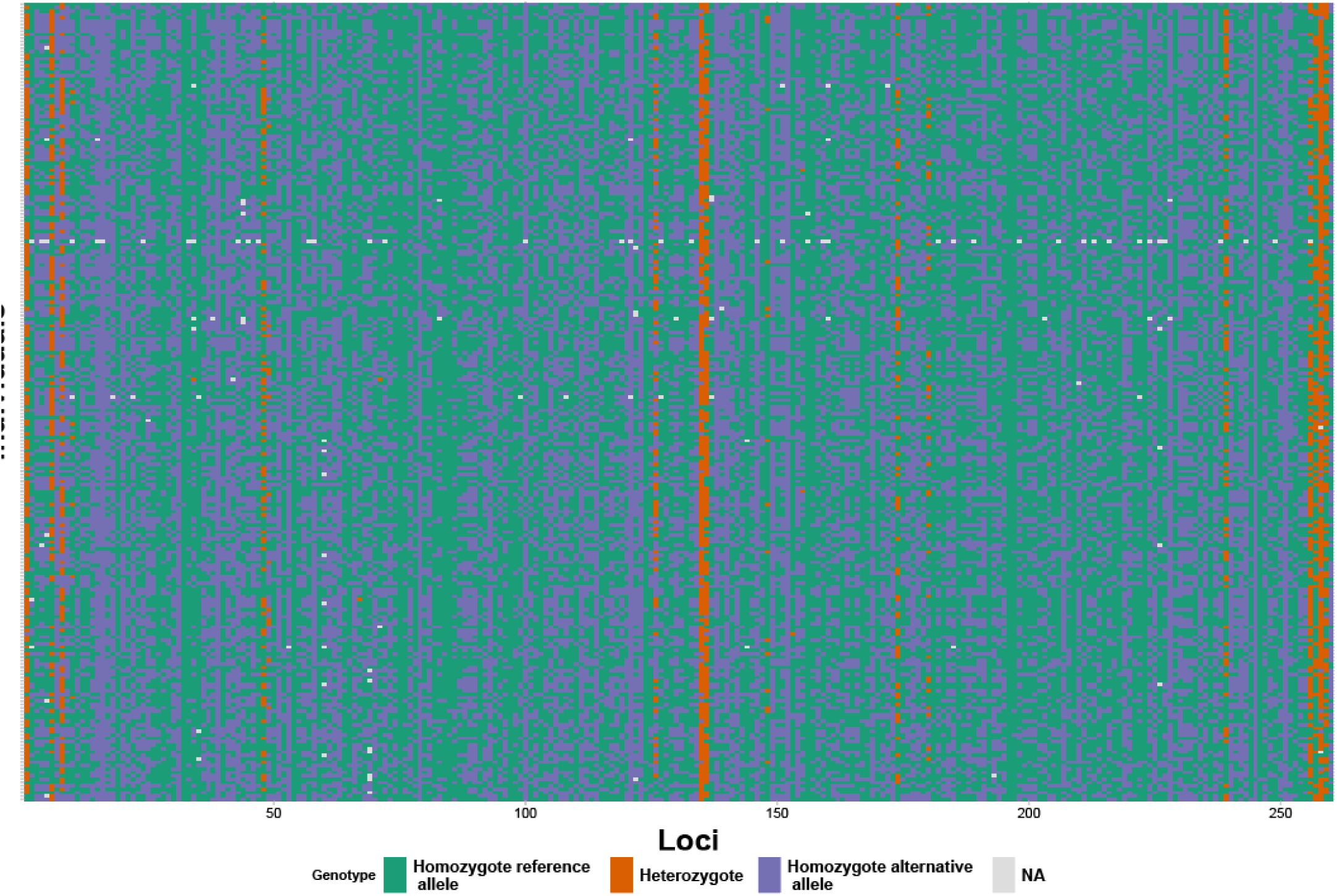
Smearplot showing genotypes of all impaternate individuals across all generations in this study, showing patterns in which loci maintain heterozygosity in parthenogenetic *Megacrania batesii* individuals. SNP loci 1 – 260 (numbered arbitrarily) are represented along the vertical axis and individuals are stacked vertically in no particular order. Colour represents genotype calls at each locus for each individual: Green represents a homozygous reference genotype, orange represents a heterozygote genotype, blue represents a homozygous alternative allele genotype, and white represents missing data.

## Notes

### Competing Interest Statement

The authors have declared no competing interest.

### Summary of Updates

An author was omitted by mistake. They have now been added to the author list.

